# Cellular substrates of functional network integration and memory in temporal lobe epilepsy

**DOI:** 10.1101/2021.01.31.428369

**Authors:** Linda Douw, Ida A. Nissen, Sophie M.D.D. Fitzsimmons, Fernando A.N. Santos, Arjan Hillebrand, Elisabeth C.W. van Straaten, Cornelis J. Stam, Philip C. De Witt Hamer, Johannes C. Baayen, Martin Klein, Jaap C. Reijneveld, Djai B. Heyer, Matthijs B. Verhoog, René Wilbers, Sarah Hunt, Huibert D. Mansvelder, Jeroen J.G. Geurts, Christiaan P.J. de Kock, Natalia A. Goriounova

**Affiliations:** Department of Anatomy and Neurosciences, Amsterdam UMC, Vrije Universiteit Amsterdam, Amsterdam Neuroscience, the Netherlands; Department of Radiology, Athinoula A. Martinos Center for Biomedical Imaging, Massachusetts General Hospital, USA; Department of Clinical Neurophysiology and MEG Center, Amsterdam UMC, Vrije Universiteit Amsterdam, Amsterdam Neuroscience, the Netherlands; Department of Neurosurgery, Amsterdam UMC, Vrije Universiteit Amsterdam, Amsterdam Neuroscience, VUmc Cancer Center Amsterdam Brain Tumor Center Amsterdam, the Netherlands; Department of Medical Psychology, Amsterdam UMC, Vrije Universiteit Amsterdam, Amsterdam Neuroscience, VUmc Cancer Center Amsterdam Brain Tumor Center Amsterdam, the Netherlands; Department of Neurology, Amsterdam UMC, Vrije Universiteit Amsterdam, Amsterdam Neuroscience, VUmc Cancer Center Amsterdam Brain Tumor Center Amsterdam, the Netherlands; Stichting Epilepsie Instellingen Nederland (SEIN), Heemstede, the Netherlands; Department of Integrative Neurophysiology, Center for Neurogenomics and Cognitive Research (CNCR), Vrije Universiteit Amsterdam, Amsterdam Neuroscience, the Netherlands; Department of Human Biology, Division of Cell Biology, Neuroscience Institute, University of Cape Town, South Africa

**Keywords:** action potential kinetics, cellular morphology, connectome, graph theory, resting-state fMRI

## Abstract

Temporal lobe epilepsy (TLE) patients are at risk of memory deficits, which have been linked to functional network disturbances, particularly of integration of the default mode network (DMN). However, the cellular substrates of functional network integration are unknown. We leverage a unique cross-scale dataset of therapy-resistant TLE patients, who underwent fMRI, MEG and/or neuropsychological testing before neurosurgery. fMRI and MEG underwent atlas-based connectivity analyses. Functional network centrality of the lateral middle temporal gyrus, part of the DMN, was used as a measure of local network integration. Subsequently, non-pathological cortical tissue from this region was used for single cell morphological and electrophysiological patch-clamp analysis, assessing integration in terms of total dendritic length and action potential rise speed. As could be hypothesized, greater network centrality related to better memory performance. Moreover, greater network centrality correlated with more integrative properties at the cellular level across patients. We conclude that individual differences in cognitively relevant functional network integration of a DMN region are mirrored by differences in cellular integrative properties of this region in TLE patients. These findings connect previously separate scales of investigation, increasing translational insight into focal pathology and large-scale network disturbances in TLE.

---

Temporal lobe epilepsy (TLE) is hallmarked by localized pathology of the temporal lobe. It is often accompanied by cognitive disturbances and particularly memory deficits (Meador 2002), which are poorly understood through local pathological markers alone. Indeed, cognition is increasingly thought to depend on the orchestrated functional dynamics taking place on a large-scale structural network of connected brain regions (Stam 2014; Bassett and Sporns 2017). These functional network dynamics can be studied using functional MRI (fMRI) and magnetoencephalography (MEG), which assess synchronized brain activity of different regions.

The most important properties of brain networks in relation to cognition are segregation and integration (Deco et al. 2015; Cohen and D’Esposito 2016; Horien et al. 2019). Segregation refers to the extent to which nodes are locally interconnected, and integration signifies the level of integrative connectivity taking place, either in globally or at a particular node. At specific brain regions, the extent of segregation and integration typically show opposite patterns (van den Heuvel and Sporns 2011; Bertolero et al. 2017): certain regions have a more segregative topological role (e.g. regions within the visual system), while others are considered mainly integrative (e.g. regions in the frontal lobe), indicating that brain regions tend to have a ‘typical’ role in the brain network that can be quantified with either segragative or integrative network measures. For example, centrality indicates the expected level of network integration occurring at any particular node, with nodes showing high integration most likely having low segregation.

Integrative connectivity of the temporal lobe, also overlapping with the default mode network (DMN (Raichle et al. 2001)), may be paramount to explain individual memory differences in TLE (DeSalvo et al. 2014; McCormick et al. 2014; Douw et al. 2015). An exemplar study explored the integrative connectivity of the hippocampal circuit with the posterior DMN (Voets et al. 2014) reporting that greater memory deficits were related to lower network integration. An open question remains whether combining different imaging and neurophysiological modalities may improve explanation of cognitive variance in these patients. Multilayer network theory provides a framework to integrate such modalities into a single network consisting of interconnected layers (De Domenico et al. 2013). Previous work suggests that multilayer functional brain network integrative measures of centrality supersede unilayer properties in explaining cognitive functioning in Alzheimer’s disease patients (Yu et al. 2017), but this approach has not been used in TLE or on combined fMRI and MEG networks.

Moreover, the cellular substrates of functional network integration are unknown, limiting our understanding of how focal cellular properties and pathology relate to large-scale network disturbances in TLE and other neurological diseases (Bassett and Bullmore 2009; Stam 2014). “Microstructure-informed connectomics” (Larivière et al. 2019) may connect these scales of measurement (Sporns 2016; van den Heuvel and Yeo 2017). In animals, more integrative structural network regions are comprised of bigger neurons with more axons (Scholtens et al. 2014). Moreover, cross-scale relations between structural brain properties covary with disease characteristics in postmortem studies of multiple sclerosis (Kiljan et al. 2019) and Alzheimer’s disease (Jonkman et al. 2020). However, the cellular substrates of functional network integration as an important correlate of cognitive impairment have been impossible to investigate *in vivo*.

We leverage a unique cohort of TLE patients undergoing functional neuroimaging and clinical neurophysiological recording as well as tissue extraction of the lateral middle temporal gyrus, a DMN region, through resective neurosurgery (Goriounova et al. 2018). We expected greater network integration to associate with more integrative cellular characteristics in terms of morphology and action potential kinetics (Poirazi et al. 2003; Eyal et al. 2014; Testa-Silva et al. 2014; Goriounova et al. 2018).

## Materials and methods

### Patients

All patients undergoing resective neurosurgery for drug-resistant epilepsy localized in the medial temporal lobe between 2009 and 2016 at Amsterdam University Medical Centers (location VUmc, Amsterdam, The Netherlands) were eligible for participation. All patients underwent temporal lobectomy, which included the lateral middle temporal gyrus. A schematic of the cross-scale analysis pipeline is provided in Figure 1.

**Figure 1.**
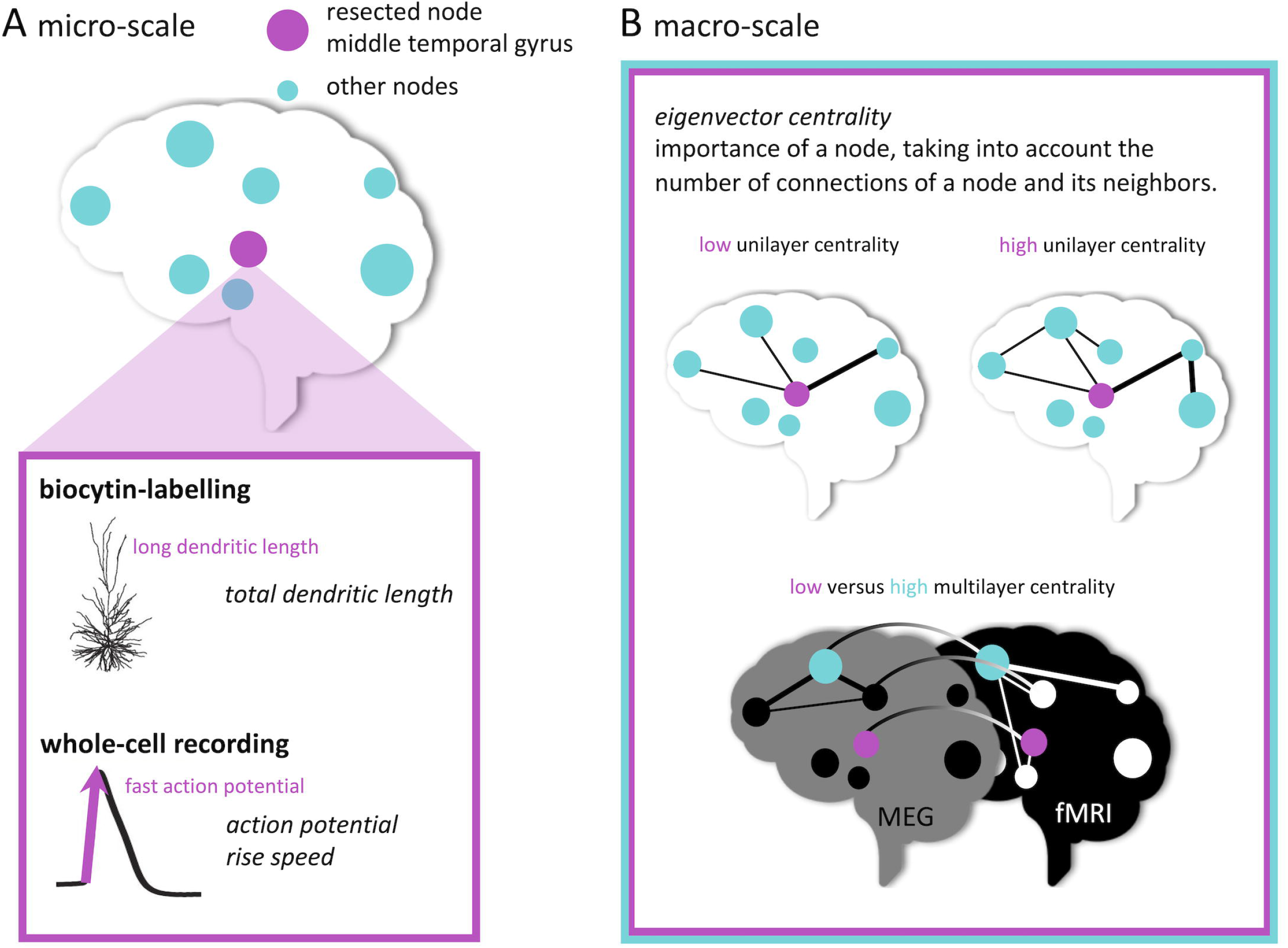
Schematic representation of multi-scale analyses In (A), cellular tissue collection from the middle temporal gyrus (pink node) is depicted. Morphological analysis and electrophysiological recordings were performed. In (B), the functional network measure of eigenvector centrality is illustrated for the unilayer (fMRI or MEG) and multilayer (combined fMRI and MEG) network analyses.

All procedures were performed with the approval of the Medical Ethical Committee of VUmc, and in accordance with Dutch license procedures and the Declaration of Helsinki. Written informed consent was provided by all subjects for data and tissue use for scientific research.

### Data Availability Statement

Patients did not consent to sharing their raw data. However, the full derived variable set used for analysis in the current work is available from GitHub (https://github.com/multinetlab-amsterdam/projects/tree/master/multiscale_integration), where our code to do so can also be found.

### Memory functioning

Patients underwent cognitive assessments during presurgical workup, as previously reported (Goriounova et al. 2018). In order to assess verbal memory functioning, the Wechsler Memory Scale (WMS) and the Dutch version of Rey’s Auditory Verbal Learning Test (RAVLT) were selected for analysis. From the WMS, the composite Verbal Memory Index was used, which has a mean value of 100 with a standard deviation of 15 in a normative, healthy population. From the RAVLT, immediate recall in terms of total correctly encoded words across five trials (15 words each) was quantified, with possible scores ranging between 0-75 words in total. Additionally, delayed recall of the 15 words was assessed after 15mins, yielding an additional outcome measure of memory retrieval. Higher values indicate better memory performance.

### Structural neuroimaging

MRI was performed on a 1.5T magnet (Siemens Sonata) and included an anatomical 3D T1-weighted MPRAGE scan (sequence parameters: TR = 2700ms, TE = 5.2ms, TI = 950ms, 1mm isotropic resolution, 176 slices). Image processing was performed using FSL5. Standard procedures were used to preprocess structural imaging: non-brain tissue was removed from the 3D T1-weighted images using the Brain Extraction Tool, and grey and white matter segmentation was performed using FAST. Non-brain tissue was removed and tissue segmentation was performed. To construct each individual’s functional brain network, the Automated Anatomical Labeling atlas was used to define 78 cortical regions. This atlas was warped from standard space to native space, and masked with each subject’s native grey matter mask.

### Functional magnetic resonance imaging

Patients underwent presurgical fMRI for language localization. Previous studies have shown that the intraindividual effect of any task or state on network topology is small in comparison to interindividual differences in network topology (Gratton et al. 2018; Kraus et al. 2021), indicating that we may use these pseudo resting-state data to investigate individual differences in network integration. Resting-state analysis has previously been used on such task data for network analysis in both healthy controls and patients (Harris et al. 2014; Krienen et al. 2014; Derks et al. 2017).

Scanning was performed using a standard echo-planar imaging sequence (TR = 2850ms, TE = 60ms, 144 volumes, 3.3mm isotropic resolution, 7min). During the scan, patients performed a language task in which nine volumes of word generation were alternated with nine volumes of rest (imagery of a landscape).

Preprocessing was performed using standard procedures (Beckmann et al. 2005), including discarding the first five volumes, motion correction, spatial smoothing, and high-pass filtering. Six regions with low signal quality (mainly orbitofrontal areas) were excluded, leaving 72 cortical regions for analysis. Additionally, ICA-AROMA was applied to minimize the impact of movement (Pruim et al. 2015). Functional images were co-registered to the anatomical scans. Time series were extracted from the centroids of all regions, after which a 72×72 connectivity matrix per subject was created using Pearson correlation coefficients. Finally, the absolute values of these correlations were used as weighted connectivity.

### Magnetoencephalography

Patients underwent resting-state MEG as part of their presurgical work-up and/or in the setting of scientific research (Nissen et al. 2017, 2018). Interictal eyes-closed recordings were acquired in supine position using a whole-head system (Elekta Neuromag Oy, Helsinki, Finland) with 306 channels inside a magnetically shielded room (Vacuumschmelze GmbH, Hanau, Germany). Data were recorded with a sampling frequency of 1250Hz, filtered online with a 410Hz anti-aliasing filter and a 0.1Hz high-pass filter. The head position relative to the sensors was recorded continuously with head-localization coils. A 3D digitizer (Fastrak, Polhemus, Colchester, VT, USA) digitized the head-localization coil positions and scalp outline (roughly 500 points) to allow surface matching of the scalp surface points with anatomical MRI.

Three eyes-closed resting-state recordings of typically 15 min each were recorded for clinical analysis of interictal epileptiform activity. Only one recording was analyzed in this study and chosen according to the following criteria with descending priority: (1) consisting of at least 5 minutes of data, (2) displaying the smallest number of artifacts as per visual inspection, and (3) being the earlier dataset of the three recordings.

Further analysis of these data has been extensively described before (Nissen et al. 2017). Offline spatial filtering of the raw data removed artifacts using the temporal extension of Signal Space Separation (tSSS) using MaxFilter software (Elekta Neuromag Oy; version 2.1). The reconstruction of neuronal sources was performed with an atlas-based beamforming approach, after which time series (virtual electrodes) for each centroid of each atlas region were reconstructed (Hillebrand et al. 2016). These time series were then filtered in the theta band (4-8Hz), because of its proven relevance for cognitive functioning in these patients (van Dellen et al. 2009; Douw et al. 2010), and to limit the number of investigated variables in this limited sample.

As a measure of functional connectivity, the phase lag index (PLI) was used. The PLI assesses the phase-relationship between two regions by quantifying the asymmetry in the distribution of instantaneous phase differences between two time series. It is robust against zero-lag phase synchronization due to volume conduction or field spread. This analysis yielded a 78×78 MEG connectivity matrix per patient.

### Network analysis

We then created minimum spanning trees (MSTs) to extract the functional backbone of each fMRI and MEG network and adequately allow for comparison between patients and modalities without having to use a subjective threshold for pairwise connectivities (Stam et al. 2014). The MST is a binary network that connects all nodes in a network without forming cycles while maximizing connectivity strength. Eigenvector centrality was then calculated per brain region. Of note, there is a plethora of network measures that measure integration, which are highly intercorrelated (Oldham et al. 2019). Instead of using multiple measures in this sample with limited statistical power, we chose to focus on eigenvector centrality as the network measure of integration. Eigenvector centrality is a spectral centrality measure, that not only takes into account the number of connections of a node, but also weighs the number of connections of its neighboring nodes (Lohmann et al. 2010).

In addition to modality-specific analysis of centrality, we also analyzed multimodal centrality, since an open question is whether combining different imaging and neurophysiological modalities may improve explanation of cognitive variance in these patients. Multilayer network theory offers an analytical framework that allows for such synergy between modalities to be captured (Mucha et al. 2010). Each layer in a multilayer network represents a network characterized by one type of connectivity. Interlayer connections link the same region across different layers. Importantly, multilayer network measures supersede summed properties of individual layers when trying to explain the behavior of other types of complex networks (Stegehuis et al. 2016). Multilayer techniques have only recently been applied to neuroscience, but show promising results towards explaining more cognitive variance than unilayer analyses in for instance Alzheimer’s disease patients (Yu et al. 2017). We therefore used the fMRI and MEG MSTs to construct a two-layer network consisting of the 72 nodes available in both modalities, where each region was connected only to itself across the two layers, forming an interconnected binary multiplex network. We then calculated multilayer eigenvector centrality per region (De Domenico et al. 2015). Ultimately, this network analysis yielded three centrality values per patient (fMRI, MEG, multilayer).

All analyses were performed using in-house Python scripts (publicly available from our GitHub page (https://github.com/multinetlab-amsterdam/projects/tree/master/multiscale_integration) in combination with the publicly available Brain Connectivity Toolbox (Rubinov and Sporns 2010) implemented in Matlab R2012a (Mathworks, Natick, MA, USA).

### Single cell electrophysiology and morphology

Tissue exclusively originated from the lateral middle temporal gyrus and was removed in order to gain access to the disease focus in deeper lying structures. In all patients, the resected neocortical tissue was not part of the epileptic focus or tumor and displayed no structural/functional abnormalities according to presurgical MRI investigation and histological analysis by an experienced pathologist. Data analysis has been extensively described before (Goriounova et al. 2018).

Upon surgical resection, the cortical tissue was immediately transferred to ice-cold artificial cerebral spinal fluid, then transported to the electrophysiology lab within 15 mins, where neocortical slices (350μm thickness) were prepared (Goriounova et al. 2018). Whole-cell patch-clamp recordings were made of layer 2 and layer 3 pyramidal neurons and action potentials (APs) were elicited by incrementing step current injections (step size 30–50 pA). Waveforms were sorted according to their instantaneous firing frequency (1/time to previous AP) and AP rise speed was defined as the peak of AP derivative (dV/dt) for all APs from all neurons from each subject.

During electrophysiological recordings, cells were loaded with biocytin through the recording pipette. After recording, slices were fixed in 4% paraformaldehyde and cells were revealed with chromogen 3,3-diaminobenzidine (DAB) tetrahydrochloride using the avidin–biotin–peroxidase method. Neurons were digitally reconstructed using Neurolucida software (Microbrightfield, Williston, VT, USA). Only neurons with virtually complete dendritic structures were included.

We then selected three representative properties pertaining to integration at the cellular scale. Larger dendrites may enable neurons to have more synaptic contacts, putatively playing a more important integrative role than neurons with smaller dendrites (Poirazi et al. 2003; Eyal et al. 2014). Larger dendrites also directly influence the speed of action potential initiation, possibly providing these cells with better temporal resolution and more efficient information transfer (Eyal et al. 2014; Testa-Silva et al. 2014; Goriounova et al. 2018). Thus, regions that act as integrators for cognitive processes may be characterized by neurons with larger dendrites and faster action potentials. We therefore extracted total dendritic length (TDL) of all basal and apical dendrites and then averaged these data (1 to 10 neurons per patient) as our first measure of cellular integration, also because this measure proved cognitive relevant in these patients before (Goriounova et al. 2018). Additionally, we selected rise speed of the 1^st^ AP, and APs fired at frequencies between 20-40 Hz for cross-scale analysis, also due to their proven relevance for cognition in this patient cohort (Goriounova et al. 2018).

### Statistical analysis

Statistical analysis was performed in Matlab R2012a (Mathworks, Natick, MA, USA).

Pairwise associations between functional network centralities and memory were tested using Spearman’s correlation coefficients. Cross-scale pairwise associations were tested using non-parametric Spearman’s correlation coefficients with bootstrapping (1 000 samples, 95% confidence intervals (CI)). When significant, robustness was explored by permuting the micro-macro pairs to create a data-specific correlation distribution (10 000 permutations) and by leave-one-out analysis. Spatial specificity of the associations was explored by correlating the cellular measure with functional network centralities of all other ipsilateral nodes in the network. The threshold for statistical significance was set at two-tailed alpha < 0.05, but we also report significant results after applying Bonferroni correction for multiple comparisons.

## Results

### Patients

In 15 of the 46 TLE patients originally included (Goriounova et al. 2018), MEG and fMRI were not available. Therefore, 31 patients (15 females) with a mean age of 33 years (± 11 years) were included for the current analysis (see Table 1 for detailed characteristics per patient).

**Table 1.**
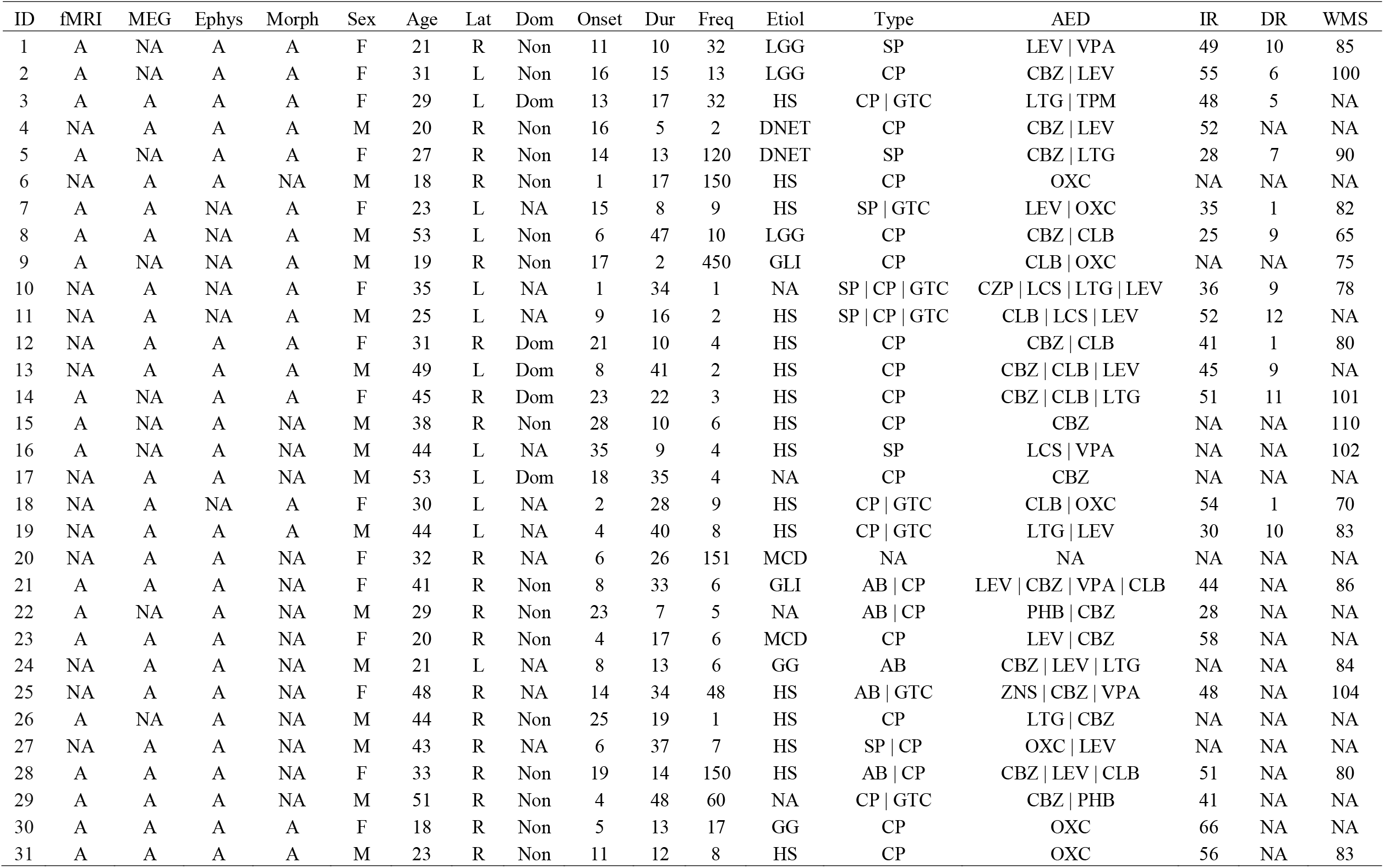
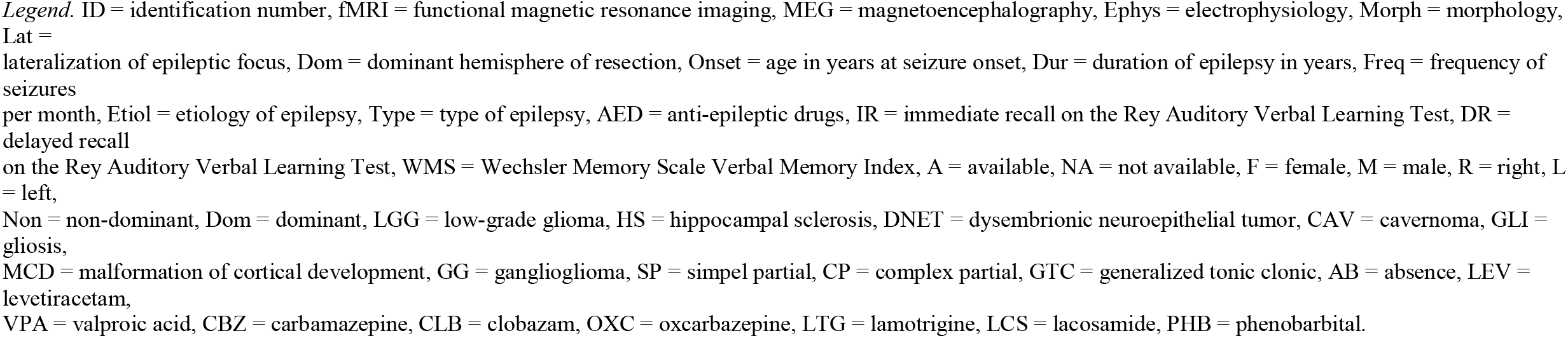
Patient characteristics

### Functional network integration and memory functioning

Firstly, we sought to confirm that functional network centrality of the lateral middle temporal gyrus, as part of the DMN, related to memory functioning in these patients. Greater fMRI network centrality was significantly related to greater RAVLT delayed recall (rho = 0.857, CI [0.373 1.000], *p* = 0.024, *n* = 7; Figure 2). Unfortunately, very few patients had complete data across these scales, rendering the remaining pairwise testing of associations underpowered.

**Figure 2.**
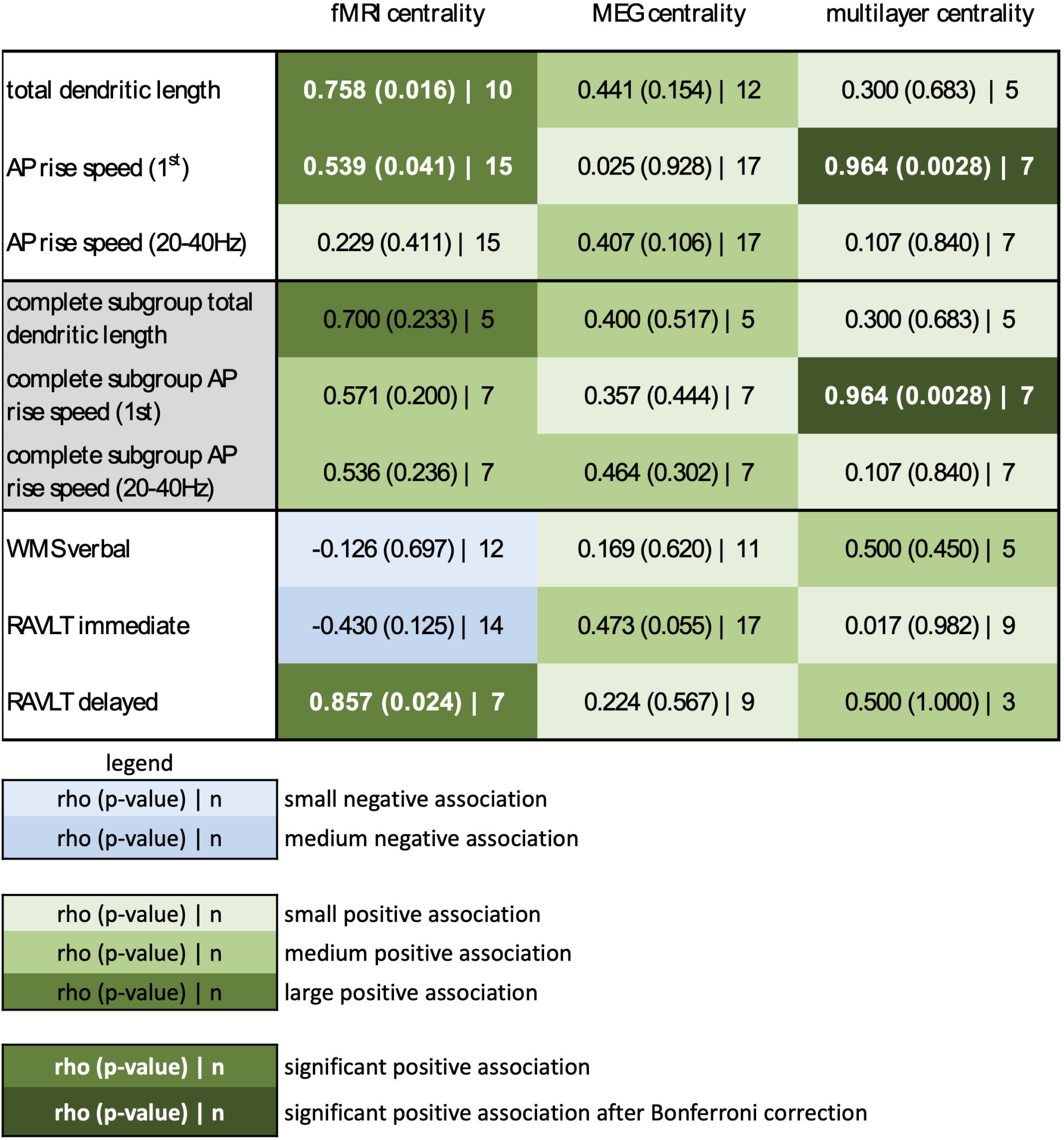
All pairwise correlations This figure shows an overview of all associations between cellular properties and functional network centrality (top three rows), as well as between functional network centrality and memory functioning (bottom three rows), using the maximum samples of patients with available data. Additionally, the middle three rows (in gray) reflect all pairwise correlations between scale-specific properties when considered in the same subgroup with complete functional network data for the morphological (*n* = 5) and electrophysiological (*n* = 7) analyses. Green elements reflect positive correlations (with rho, *p* and *n* in text), ranging from small correlations (rho < 0.4, light green) to medium correlations (0.4 < rho < 0.6, medium green) to large correlations (rho > 0.6, dark green). White, bold text indicates statistical significance (*P* < 0.05) and very dark green elements with white, bold text indicates statistical significance after Bonferroni correction for multiple (nine) comparisons. Blue elements reflect negative correlations in the same way. fMRI = functional Magnetic Resonance Imaging, MEG = magnetoencephalography, AP = action potential, WMS = Wechsler Memory Scale, RAVLT immediate = Rey Auditory Verbal Learning Test immediate recall, RAVLT immediate = Rey Auditory Verbal Learning Test delayed recall. In (B), we only select patients with both functional modalities and either of the cellular measures available, namely five patients with total dendritic length and seven patients with action potential rise speed. The legend is identical between (A) and (B).

### Cross-scale correlations of integrative properties

We then asked whether greater functional network centrality of the resected region went hand in hand with longer TDLs and faster APs, hypothetically signs of greater integrative potential at the cellular level (Figure 2). Longer TDL was significantly related to greater fMRI network centrality (rho = 0.758, CI [0.115 0.975], *p* = 0.016, *n* = 10; Figure 3A). This association remained significant when creating a sample-specific distribution of the correlation through permutation analysis (rho cut-off = 0.636, Figure 3B left panel). We then performed leave-one-out analyses, where we iteratively excluded a single patient from the correlation analysis to see whether this result was driven by individual data point. This yielded significant results in 7 of 10 analyses (Figure 3B middle panel). Finally, we explored whether TDL of the resected region also correlated with network centrality of other cortical regions. This analysis revealed a significant negative correlation between TDL of the resected region and fMRI network centrality of the ipsilateral paracentral lobule (rho = -0.721, *p* = 0.024, Figure 3B right panel), although this result did not survive correction for the 36 correlations tested.

**Figure 3.**
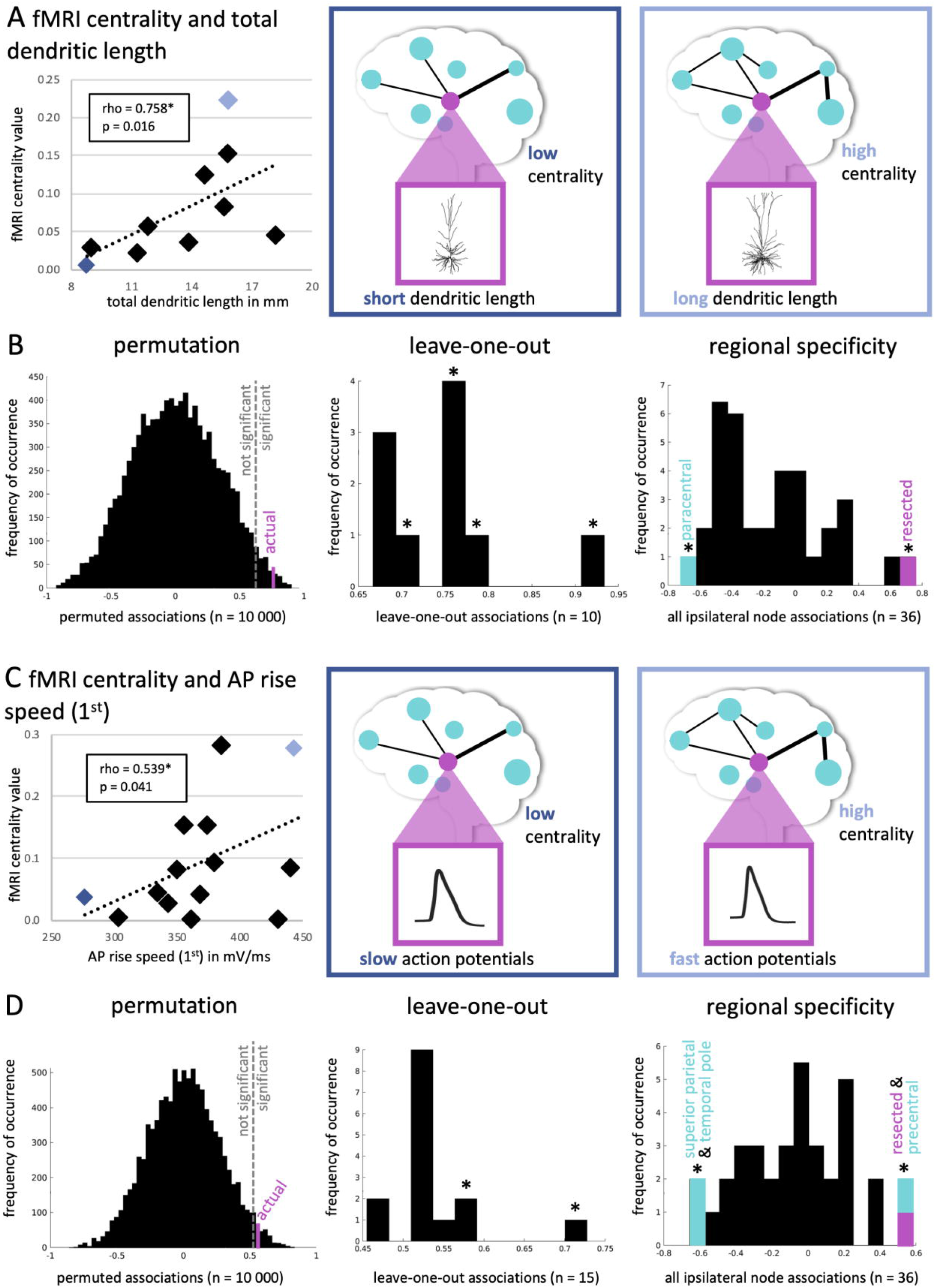
Significant pairwise associations between cellular properties and functional network centrality (A) Displays a scatter plot of the significantly positive correlation between total dendritic length (in mm) and functional Magnetic Resonance Imaging (fMRI) centrality. The dark blue diamond in the plot represents a single patient, who has low centrality and short dendritic length (schematically depicted in the middle panel), while the light blue diamond represents a patient with high centrality and long dendritic length (right panel). In (B), the analyses of the robustness of these results are displayed. The left panel shows the distribution of permuted correlations (10 000 permutations), with the actual (pink) association being smaller than the alpha = 0.05 threshold indicated by the dotted line. The middle panel displays the 10 leave-one-out associations in addition to the real correlation, yielding a significant correlation in 7 of 10 analyses as indicated by the asterisks. The right panel displays all correlations between ipsilateral functional network centrality values (*n* = 36 regions) and total dendritic length of the resected area. The reported positive association (with the resected region) in pink as well as the negative correlation between paracentral centrality and total dendritic length are significant. In (C), a scatter plot of the significantly positive correlation between action potential (AP) rise speed (1^st^) and fMRI centrality is shown. In (D), the analyses of the robustness of these results are displayed in the same manner as in (B).

Furthermore, faster AP rise speed (1^st^) was significantly related to greater fMRI network centrality (rho = 0.539, CI [0.042 0.868], *p* = 0.041, *n* = 15; Figure 3C/D). This finding remained in permutation testing (rho cut-off = 0.525), but was mostly non-significant in leave-one-out analyses. Spatially, AP rise speed (1^st^) of the resected area was also significantly associated with fMRI network centrality of the precentral region (rho = 0.582, *p* = 0.025). Moreover, there were significant negative relationships with fMRI network centrality of the superior parietal region (rho = -0.657, *p* = 0.010) and temporal pole (rho = -0.611, *p* = 0.018), although these correlations did not survive correction for multiple comparisons.

The only pairwise correlation that survived correction for the nine pairwise tests performed between cellular and functional network characteristics of the resected region was the significant positive association between AP rise speed (1^st^) and multilayer network centrality (rho = 0.964, CI [0.698 1.000], *p* < 0.001, *n* = 7; Figure 4A). Moreover, this association was robust in permutation and leave-one-out analyses, and was spatially specific, also when not correcting for multiple comparisons.

**Figure 4.**
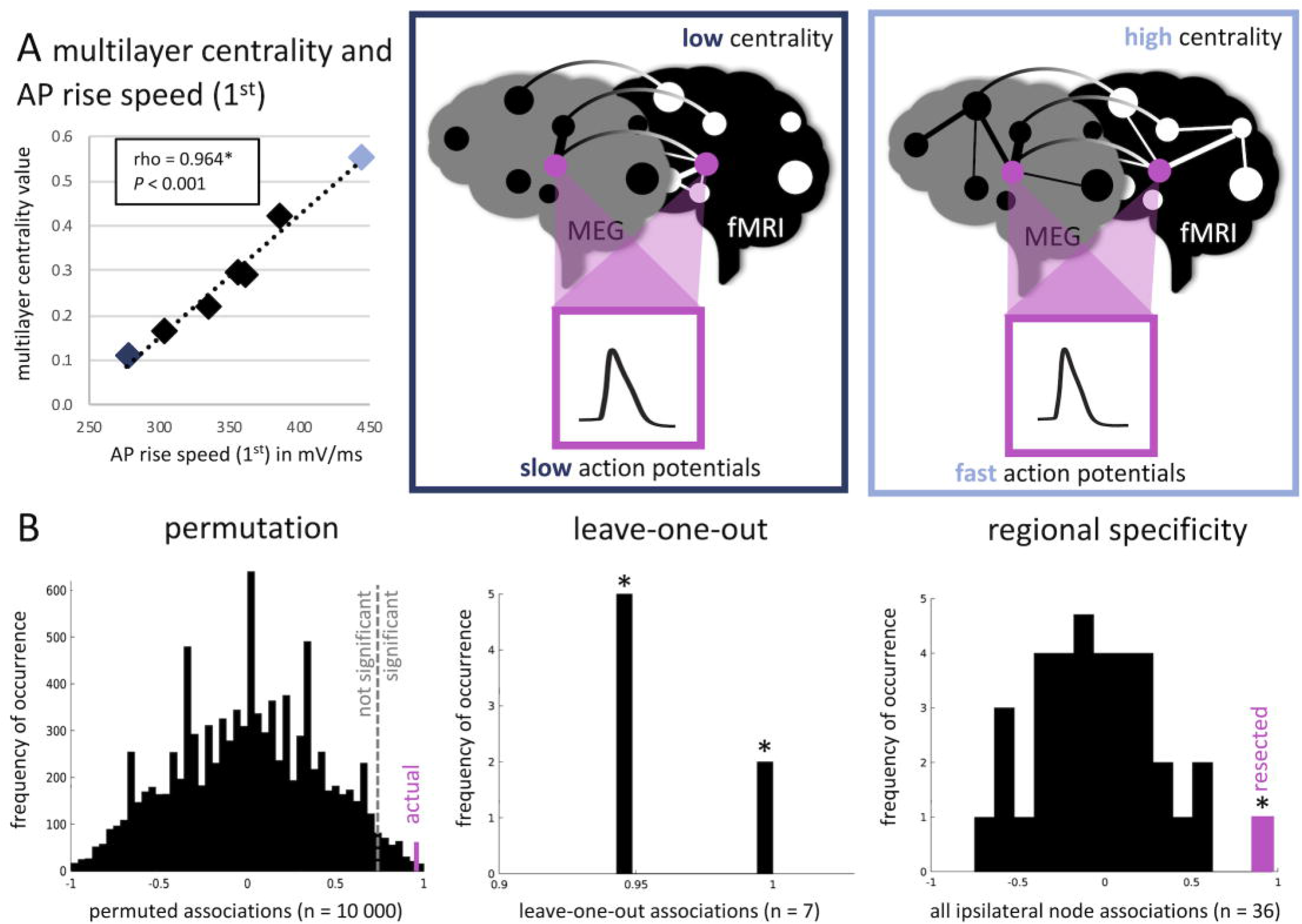
Significant pairwise association between cellular properties and functional network centrality after correction for multiple comparisons (A) Displays a scatter plot of the significantly positive correlation between action potential (AP) rise speed (1^st^) and multilayer centrality, which is the only pairwise association significant after Bonferroni correction for multiple comparisons. The dark blue diamond in the plot represents a single patient, who has low centrality and slow action potentials (schematically depicted in the middle panel), while the light blue diamond represents a patient with high centrality and fast action potentials (right panel). In (B), the analyses of the robustness of these results are displayed. The left panel shows the distribution of permuted correlations (10 000 permutations), with the actual (pink) association being smaller than the alpha = 0.05 threshold indicated by the dotted line. The middle panel displays the 7 leave-one-out associations in addition to the real correlation, yielding a significant correlation in all analyses as indicated by the asterisks. The right panel displays all correlations between ipsilateral network centrality values (*n* = 36 regions) and AP rise speed (1^st^) of the resected area. The reported positive association (with the resected region) in pink is the only significant correlation.

Since different numbers of patients were available for each pairwise correlation depending on modalities available, we also investigated whether the small subset of patients with complete data showed the same patterns of correlation as the larger but heterogeneous samples. Indeed, Figure 2 also shows the associations of both unilayer and multilayer network indices with cellular measures in the same samples of 5 patients (for TDL) and 7 patients (for AP rise speeds). As was the case for the mixed samples, only multilayer network centrality and AP rise speed (1^st^) were significantly correlated in this complete dataset, indicating that patient selection differences between the above described subsamples did not confound the reported pairwise correlations.

## Discussion

We report on a unique cohort of TLE patients with data spanning multiple scales of investigation, in order to explore the cellular substrates of cognitively relevant functional network integration of a DMN region. As expected (McCormick et al. 2014; Voets et al. 2014; Douw et al. 2015), greater functional network integration of the lateral middle temporal gyrus related to better memory functioning in these patients. Moreover, greater network centrality of this brain region correlated with signs of greater cellular integration: patients with longer dendrites and faster action potentials also displayed more integrative functional network profiles.

Our findings signify that brain organization in terms of integrative propensity is preserved across scales of measurement in TLE patients. Functional network centrality of the investigated DMN region correlated with neuronal morphology in terms of total dendritic length of its constituent neurons. In previous work, we showed that pyramidal neurons of patients with higher intelligence in the related cohort had larger, more complex dendrites (Goriounova et al. 2018). Larger dendrites would enable neurons to have more synaptic contacts, putatively playing a more important integrative role than neurons with smaller dendrites (Poirazi et al. 2003; Eyal et al. 2014). Greater dendritic length and axonal density of neurons at particular locations have previously been linked to structural network integration as measured with diffusion MRI: more integratively connected network regions tended to have bigger neurons with more axons, while locally connected network regions were made up of smaller neurons that were connected with lower axonal density (Scholtens et al. 2014; Kiljan et al. 2019). The current results suggest that functional network integration also mirrors more integrative neuronal morphology, at least in the lateral middle temporal gyrus as part of the DMN.

Functional network centrality also related to action potential kinetics: more integrative functional network topology related to faster action potential rise speeds. We previously found that patients with higher intelligence had faster action potentials during high frequency activity, in addition to the abovementioned longer total dendritic lengths (Goriounova et al. 2018). Of note, larger dendrites also directly influence the speed of action potential initiation, putatively offering more integrative power (Eyal et al. 2014; Testa-Silva et al. 2014; Goriounova et al. 2018). Our current results put these local indicators of integration into a cross-scale network perspective, as we report on associations with functional network centrality. Our findings support a scale-free view on these type of brain properties, such that within-region cellular integrative properties are reflected by between-region functional network integration, which is of particular relevance towards understanding (cognitive deficits in) TLE.

Of note, the functional network correlates of local action potential kinetics became particularly evident when taking both functional modalities (fMRI and MEG) into account through multilayer network theory. This finding suggests that multimodal functional centrality may capture cellular brain properties better than either of the two modalities alone, which would be in line with other multilayer brain network studies (Battiston et al. 2017; De Domenico 2017). It should, however, be noted that the lack of robust significance for unilayer network centralities as well as the significance of multilayer network centrality may reflect limited statistical power in small subsamples.

Based on our findings, richer hypotheses may be formulated on the relationships between local cellular organization and functionality on the one hand, and large-scale network topology on the other hand in TLE. Next steps may include investigation of cellular and network properties of pathological brain regions in addition to the non-pathological area of the DMN assessed in this study. This would bridge the gap between focal TLE pathology and its widespread cross-scale effects on the brain, ultimately pertaining to cognitive impairment. Such studies are of particular interest, since our current finding of cross-scale preservation of integrative propensity in a non-pathological brain region may not necessarily be specific to TLE: previous work in macaques and postmortem donors also report on such correlations (Scholtens et al. 2014; Kiljan et al. 2019; Jonkman et al. 2020), raising the question whether cross-scale integration is a basic organizational principle conserved across species to begin with. By also involving focal pathological cellular properties (and their large-scale network counterparts), disease-specific processes may be disentangled from such fundamental principles.

Several limitations of this study should be noted. First and foremost, the unique nature of this multi-scale dataset meant that only a small sample was available, also precluding subgroup analysis or exploration of confounders that might have affected the current results, e.g. lateralization of the epileptogenic zone, sex, and age. It was also impossible to use a control group for this analysis: healthy individuals can obviously not be subjected to the cellular measurements we report on. Another limitation is the spatial resolution of matching between the cellular and functional network analyses: while certainty about the location of origin of the resected tissue was in the order of millimeters, the atlas region used to reflect the tissue location spans several centimeters. We chose this atlas as it has been used successfully by our group in previous studies with comparable patient populations and thus has proven cognitive and cellular relevance (van Dellen et al. 2012, 2014; Carbo et al. 2017), but future studies may aim to match tissue locations with increased spatial accuracy.

In conclusion, we show that individual differences in functional network integration of a DMN region relate to cellular morphology and action potential kinetics. These results underline the translational nature of individual differences in brain properties between TLE patients, which has clinical relevance in terms of memory functioning. Ultimately, such “microstructure-informed connectomics” (Larivière et al. 2019) may lead the way towards better understanding and treatment of neurological disease in general, and memory functioning in TLE patients specifically.

## Acknowledgements

We thank Lucas Breedt for his help in testing the multilayer code implementation. Part of the data collection was funded by the Nederlands Epilepsie Fonds (NEF 08-08, 09-09 for Dr. Douw and Dr. Reijneveld), the Dutch Organization for Scientific Research (NWO Veni 016.146.086 and NWO Vidi 198.015 for Dr. Douw), Society in Science (Branco Weiss Fellowship for Dr. Douw). Dr. Douw, Dr. Goriounova and Dr. De Kock received funding from Amsterdam Neuroscience for this work. Dr. Goriounova received funding for this work from the Dutch Organization for Scientific Research (NWO Veni, 016.Veni.171.017) and from Amsterdam Neuroscience. Dr. Nissen and Dr. Hillebrand were funded by the Nederlands Epilepsie Fonds (NEF 14-16) and the Netherlands Organisation for Health Research and Development (ZonMW grant 95105006). Prof. Mansvelder received funding from the European Union’s Horizon 2020 Framework Programme for Research and Innovation (grant 785907 for Human Brain Project SGA2 and SGA3). These funding bodies were not involved in the analysis or interpretation of results or manuscript writing.

